# A flaw in using pre-trained pLLMs in protein-protein interaction inference models

**DOI:** 10.1101/2025.04.21.649858

**Authors:** Joseph Szymborski, Amin Emad

## Abstract

With the growing pervasiveness of pre-trained protein large language models (pLLMs), pLLM-based methods are increasingly being put forward for the protein-protein interaction (PPI) inference task. Here, we identify and confirm that existing pre-trained pLLMs are a source of data leakage for the downstream PPI task. We characterize the extent of the data leakage problem by training and comparing small and efficient pLLMs on a dataset that controls for data leakage (“strict”) with one that does not (“non-strict”). While data leakage from pre-trained pLLMs cause measurable inflation of testing scores, we find that this does not necessarily extend to other, non-paired biological tasks such as protein keyword annotation. Further, we find no connection between the context-lengths of pLLMs and the performance of pLLM-based PPI inference methods on proteins with sequence lengths that surpass it. Furthermore, we show that pLLM-based and non-pLLM-based models fail to generalize in tasks such as prediction of the human-SARS-CoV-2 PPIs or the effect of point mutations on binding-affinities. This study demonstrates the importance of extending existing protocols for the evaluation of pLLM-based models applied to paired biological datasets and identifies areas of weakness of current pLLM models.

## Introduction

Protein large language models (pLLMs) have become a driving force in pushing new frontiers for biological tasks including annotating protein domains and families [1], predicting microbial virulence [2], and of most interest to the authors, protein-protein interaction (PPI) inference [3]. Today, the rich and compact vector representations of proteins that pLLMs offer are an attractive set of features for researchers constructing PPI inference models. High-scoring pLLM-based PPI prediction models may, however, elicit skepticism in members of the community familiar with the pernicious challenge of data leakage and generalization that often plague models inferring PPIs. What follows is an in-depth study of the suitability and robustness of pre-trained pLLMs for downstream PPI inference tasks which we have undertaken to determine whether fears of data leakage are founded.

Unravelling the interactions between proteins is key to unlocking the inner workings of fundamental biological processes and has played a foundational role in our understanding of molecular biology and disease [4]. Protein-protein interactions (PPIs) are traditionally studied through wet-lab methods like co-immunoprecipitation (Co-IP), pull-down assays, and high-throughput techniques such as yeast two-hybrid (Y2H) systems or mass spectrometry approaches (*e*.*g*., CF-MS, XL-MS) [5, 6, 7]. While these methods provide reliable results, they are labor-intensive, time-consuming, and often low-throughput. To address these limitations, computational models have emerged as a cost-effective alternative, offering rapid PPI inference with minimal resource investment. It is in pursuit of lowering the high cost of impactful biological discoveries that researchers seek to build ever-more-accurate PPI inference models.

The process of pre-training pLLMs is commonly conducted on datasets such as UniRef which are constructed without regard to the evaluation of downstream tasks, like PPI inference. As a consequence, most if not all proteins which are present in the testing dataset of the downstream PPI task are certain to appear in the dataset used to pre-train pLLMs, which may lead to data leakage. When used upstream of PPI inference tasks, pre-trained pLLM encoders almost always constitute the vast majority of parameters in the network, exacerbating the risk that pre-trained pLLMs can cause PPI inference tasks to leak data. It is this maximalist approach to training data, driven by the large number of network parameters, that is at the heart of the concern for data leakage.

Over a decade ago in a letter to Nature Methods, Park and Marcotte revealed a critical flaw regarding the methods used to train and evaluate PPI inference models that results in dramatically inflated performance metrics [8]. Namely, when test protein pairs share constituent proteins with training data, artificially appear to achieve high Area Under the Receiver Operating Characteristic curve (AUROC) scores despite earning scores matching random-chance methods on testing sets with held-out proteins. Despite the advent of deep learning, this issue persists —recent work by Bernett et al. (2024) shows leading methods still fail without partial protein overlap in train/test sets, underscoring ongoing methodological weaknesses [9].

Further complicating matters is homology-driven leakage, where proteins share near-identical sequences due to evolutionary conservation [10]. Even if test and training sets avoid direct protein overlaps by name, sequence similarity (e.g., from conserved domains) allows information transfer. Thus, robust PPI evaluation requires stringent checks for both explicit protein overlap and amino acid sequence identity across datasets.

In light of previous findings of the pervasive nature of sources of data leakage in PPI inference models, it stands to reason that pLLMs which are pre-trained on datasets which include the testing proteins of downstream PPI tasks are a potential source of data leakage which warrant further study. An evident mitigation for the risk of data leakage in pLLMs is to pre-train pLLMs on datasets which exclude proteins which are found in the test dataset used to evaluate downstream PPI tasks. This represents an increase of many orders of magnitude to both training time and costs. If it is discovered that pre-trained pLLMs are inflating evaluation metrics, it could demand of us much more compute and labour than previously thought to repurpose pLLMs for certain downstream tasks. As such, we believe it is of utmost importance for the extent (if any) of data leakage caused by pre-trained pLLMs to be fully characterized and understood.

By pre-training a series of pLLMs on datasets which either included or excluded proteins found in downstream PPI testing pairs, we were able to show experimentally that there is indeed a degree of data leakage or loss of generalizability attributable to certain pre-trained pLLMs. Namely, pre-trained pLLMs whose training datasets are composed of proteins included in the testing dataset of downstream PPI tasks resulted in inflated evaluation metrics. Notably, while this was true for the PPI task, no such distortion in evaluation metrics was found for a different downstream task (protein keyword annotation task).

We further go on to characterize the performance of fine-tuned pLLMs on the downstream PPI inference task and the effects of pLLMs’ context-length on accurately inferring interactions between proteins with long amino acid sequences. We further identify limitations in the ability of PPI inference methods, both those based on pLLMs and those which are not, to make predictions on very out-of-distribution proteins such as those found in SARS-CoV-2, as well as their insensitivity to small-but-important changes in amino acid sequences which alter binding affinity.

## Results

### Inclusion of PPI testing proteins in the training of pLLMs inflates performance, suggesting data leakage

Prior to assessing the effect of data leakage, we evaluated the PPI prediction performance using embeddings obtained from five state-of-the-art (SOTA) pLLMs [11, 12, 13, 14], trained irrespective of the PPI testing set (Methods, Supplementary Table 1). All PPI training, validation, and testing datasets were processed according to Park & Marcotte’s “C3” definition [8] meaning the PPI training dataset did not contain interactions between proteins found in the held-out testing set (Methods). Our results (Table 1) revealed that these pLLM-based models enjoy significantly better performance compared to non-pLLM-based PPI prediction models [15, 16, 17, 18, 19].

**Table 1.** Performance of pLLM-based and non-pLLM-based PPI inference models.

| Encoder | pLLM-based? | AUROC |  | AP |  | MCC@50% |  |
| --- | --- | --- | --- | --- | --- | --- | --- |
|  |  | Mean | Std. Dev. | Mean | Std. Dev. | Mean | Std. Dev. |
| ProtT5-XL | ✓ | 0.879 | 0.00589 | 0.867 | 0.00877 | 0.580 | 0.00987 |
| ESM2-650M | ✓ | 0.866 | 0.00425 | 0.851 | 0.00266 | 0.556 | 0.00965 |
| ProtBERT | ✓ | 0.842 | 0.00583 | 0.836 | 0.00464 | 0.505 | 0.00859 |
| SqueezeProt-U50 | ✓ | 0.825 | 0.00251 | 0.804 | 0.00633 | 0.484 | 0.00381 |
| ProSE | ✓ | 0.815 | 0.00524 | 0.799 | 0.00455 | 0.467 | 0.0144 |
| ProteinBERT | ✓ | 0.810 | 0.00742 | 0.785 | 0.00801 | 0.462 | 0.0168 |
| RAPPPID | ✗ | 0.728 | 0.0225 | 0.714 | 0.0146 | 0.303 | 0.0369 |
| D-SCRIPT | ✗ | 0.617 | 0.120 | 0.613 | 0.0816 | 0.143 | 0.143 |
| Richoux *et al.* | ✗ | 0.602 | 0.00662 | 0.607 | 0.00385 | 0.155 | 0.0131 |
| SPRINT | ✗ | 0.590 | 0.0590 | 0.632 | 0.0612 | 0.116 | 0.0907 |
| PIPR | ✗ | 0.483 | 0.00484 | 0.484 | 0.00251 | -0.0233 | 0.00734 |

AUROC: Area under the Receiver Operating Characteristic.
AP: Average Precision.
MCC@50%: Matthew’s Correlation Coefficient at a 50% threshold.
Std. Dev.: Standard Deviation.
Means and Standard Deviations are calculated for three random training/testing splits ( $n = 3$ ).

Next, we set out to establish whether data leakage is introduced into downstream PPI inference models by upstream pre-trained pLLMs. Since re-training existing SOTA pLLMs multiple times, as is required by this experiment, is prohibitively costly in terms of computational resources, time, and energy, we trained a new model we called “SqueezeProt”, an efficient and tractable pLLM that utilizes the SqueezeBERT architecture [20]. This model substitutes fully-connected operations in BERT attention mechanism for grouped convolutions (Fig. 1a, Methods) [21]. To establish the suitability of this architecture for our purposes, we first trained it on the sequences of all proteins in the UniRef50 dataset, a dataset used by multiple SOTA pLLMs. When its pre-trained embeddings were used to predict PPIs, this model (henceforth “SqueezeProt-U50”) performed comparably to other SOTA pLLMs using the same PPI prediction approach (see Methods). Notably, however, it achieved an inference speed of approximately five times faster than the fastest SOTA pLLM above (Fig. 1b, Table 1, Supplementary Table 1).

**Figure 1.**
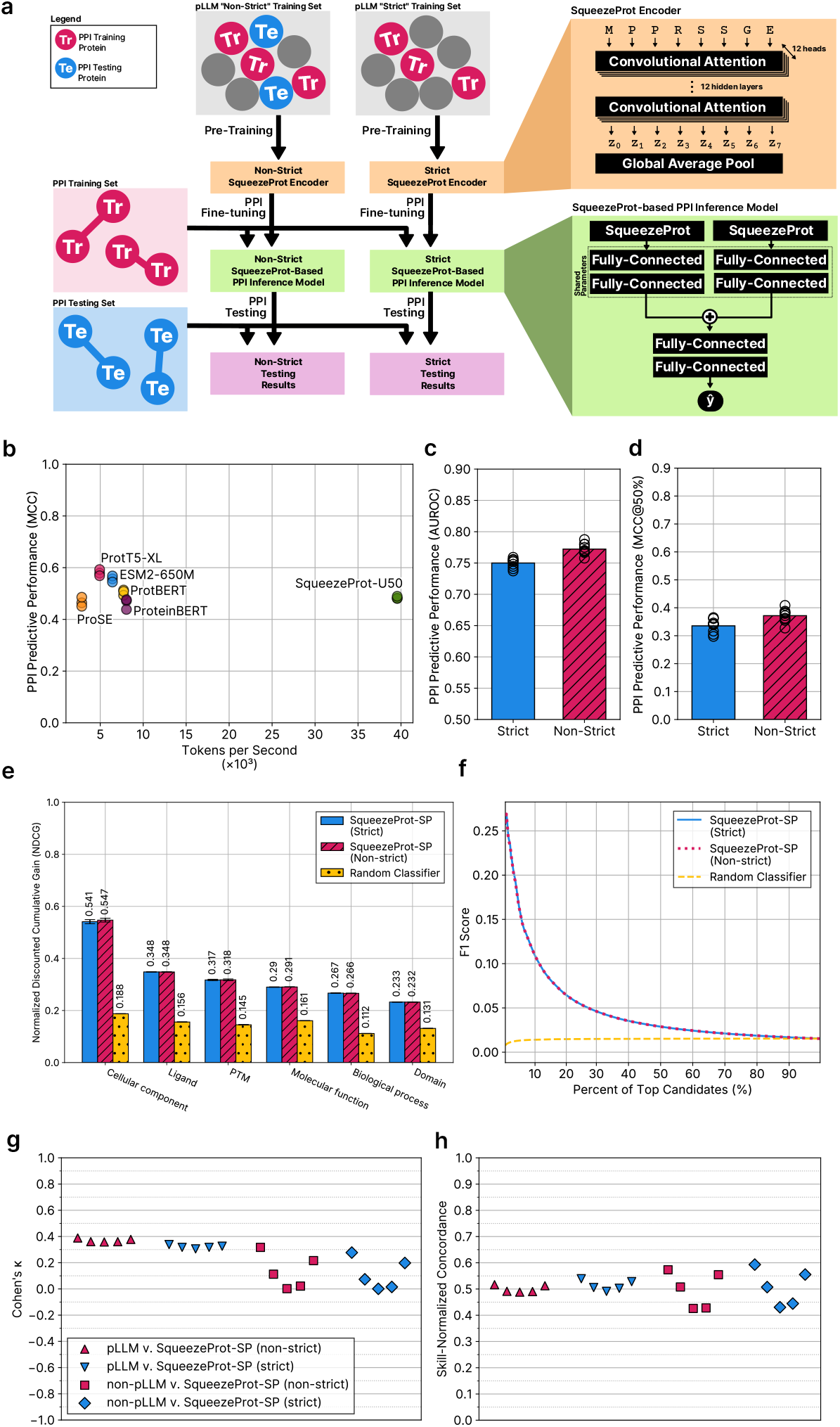
The performance of pre-trained pLLMs on downstream PPI tasks are impacted by whether or not the pLLM was trained on a strict dataset. **a**, Overview of the experimental set-up for evaluating potential changes in the PPI predictive performance due to data leakage. Non-strict and strict SqueezeProt-SP pLLMs were pre-trained using datasets that include and exclude, respectively, proteins found in the downstream PPI testing set. Both models were then fine-tuned and tested on a PPI dataset (following C3 split) using a twin neural network architecture. **b**, The predictive performance of pLLM-based PPI inference methods (measured by MCC@50%) as a function of inference speed (measured by tokens per second). Each marker represents the results of one of three models trained with different random seeds. SqueezeProt-U50 achieved comparable performance to leading pLLMs while achieving approximately 5× faster inference speed compared to fastest other pLLM. **c-d**, The mean predictive performance of the PPI inference models based on strict (blue) and non-strict (red, hatched) SqueezeProt-SP pLLM encoders, as measured by AUROC **(c)** and MCC@50% **(d)**. The mean AUROC of the strict and non-strict models is 0.750 (±0.00674) and 0.772 (±0.00966) respectively, and the mean MCC@50% of the strict and non-strict models is 0.339 (±0.0232) and 0.370 (±0.0241). Mean and standard deviation are calculated for the results of ten models trained identically but for different random seeds used to initialize weights and sample training examples. Markers indicate the performance of each of the ten models. **e-f**, Performance of models for assigning UniProt Keywords to proteins given their sequence. Models are based on strict and non-strict SqueezeProt-SP variants. **e**, The per-category testing performance, as measured by normalized discounted cumulative gain (NDCG), of models based on strict (solid blue) and non-strict (hatched red) variants of SqueezeProt-SP as compared to a random classifier (spotted yellow). The average performance of ten downstream keyword models trained with ten random seeds is reported, with error bars reflecting the standard deviation. Relevant keyword categories with more than 15 keywords are shown here. **f**, Overall testing performance, as measured by *F*_1_ score, among the top ranked candidates of strict (solid blue) and non-strict (dotted red) pLLM-based keyword models. The performance of a random classifier (dashed yellow) is also included for comparison. **g-h**, Swarm plots of the concordance between pLLM, non-pLLM, and the SqueezeProt-SP variants as measured by Cohen’s *κ* coefficient **(g)** and our skill-normalized concordance measure **(h)**. The concordance between the non-strict SqueezeProt-SP variant (red) and both pLLM (upward triangle, *n* = 5) and non-pLLM (square, *n* = 5) is reported. Additionally, concordance between the strict SqueezeProt-SP variant (blue) and both pLLM (downward triangle, *n* = 5) and non-pLLM (diamond, *n* = 5) is reported.

To further reduce the computational and time cost associated with these experiments, we focused on two smaller datasets formed from more than half a million amino acid sequences of “reviewed” proteins catalogued by the UniProtKB database (*i*.*e*., SWISS-PROT or SP) [22]. One variant was trained on a “strict” dataset in which proteins found in the PPI testing and validation sets (*n* = 2, 780) or those that shared more than 50% sequence identity with them were excluded. Another variant was trained on a “non-strict” dataset, which included proteins excluded from the strict dataset (Methods, Fig. 1 a). The two datasets above were of similar size. Pre-computed embeddings of both variants were used to train PPI inference models with identical architectures (Fig. 1a, Methods). When evaluated on the C3 PPI testing set, switching from strict to non-strict variant resulted in a noticeable increase in PPI predictive performance based on various metrics (e.g., a 9.14% increase in mean Matthews correlation coefficient (MCC)) (Fig. 1c-d, Supplementary Table 2, Supplementary Figure 1). Since the two PPI models were identical but for the data used in the pre-trained pLLM upon which they were based, we can conclude that the difference is due to data leakage in the non-strict pre-trained pLLM.

### Extent of data leakage of pLLM-based models depends on downstream tasks

Next, we sought to understand whether this data leakage extends to other protein-related tasks that do not rely on paired protein data. We chose the task of classifying proteins by the keywords assigned to them by the UniProt database [22]. These keywords belong to a controlled vocabulary, with a total of 1,119 terms assigned to 10 different categories ranging from “biological process” to “post-translational modification”. We framed this task as a multi-label classification problem and developed a fully-connected neural network which accepts as input pLLM protein embeddings and outputs a vector of probabilities, with each element corresponding to the probability of a keyword (Methods).

We observed no appreciable difference in model performance between instances that used the strict variant and its non-strict counterpart, as measured by normalized discounted cumulative gain (NDCG), Label Ranking Average Precision (LRAP), Coverage Error, and Top-3 and Top-10 Accuracy (Fig. 1e, Supplementary Table 3) [23, 24]. Similarly, when only considering the top *k* candidates ranked by models based on the strict and non-strict pLLMs, no distinction in *F*_1_ score between the two was observed for all valid values of *k* (Fig. 1f). However, a small difference was observed when measuring the accuracy and precision scores of the top *k* candidates (Supplementary Figure 2). These results suggest that the severity of data leakage in downstream tasks depends on the specific nature of each task.

**Figure 2.**
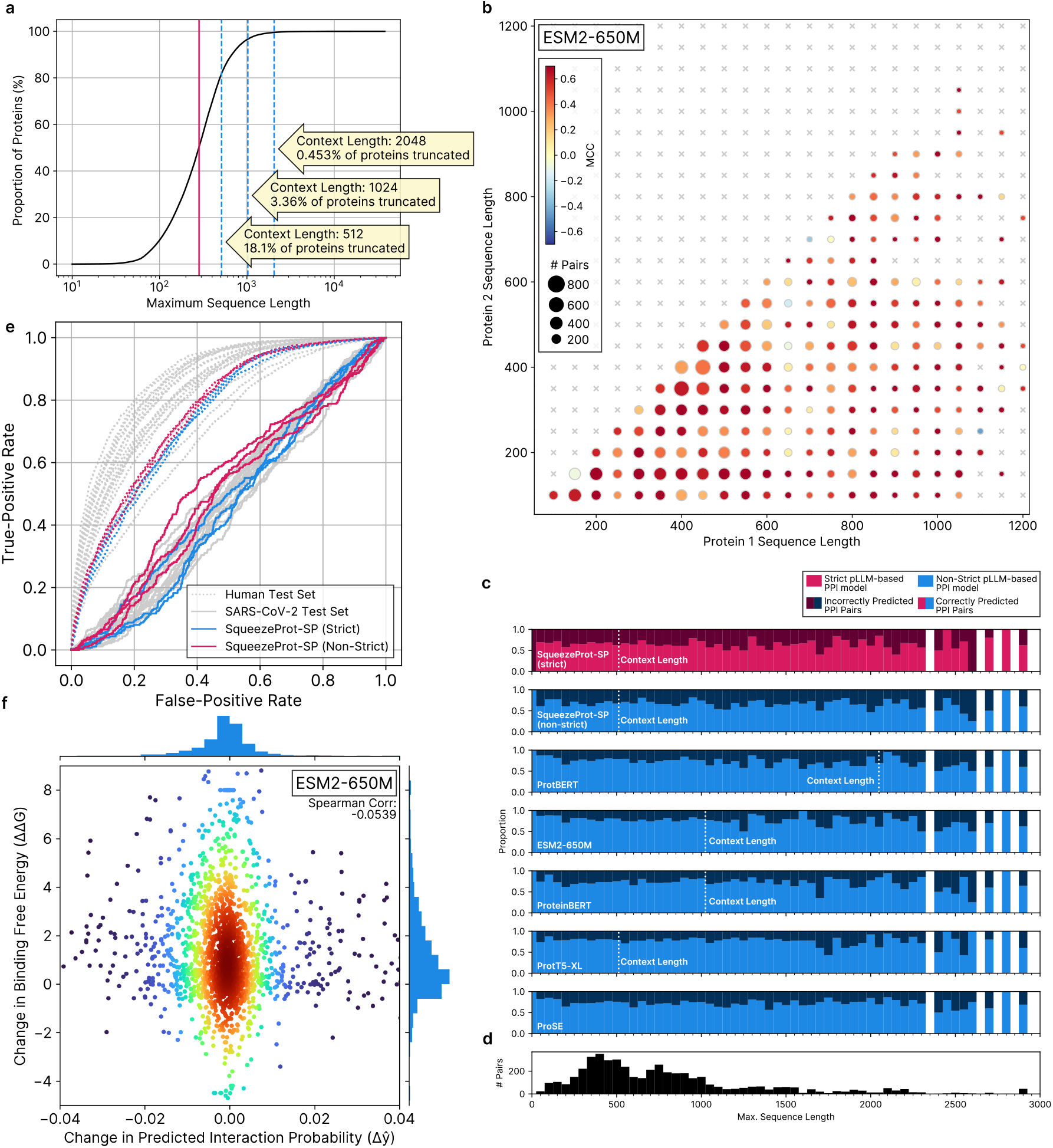
Accuracy of pLLM-based PPI inference methods is not significantly impacted by sequence length, but out-of-distribution PPIs and mutation sensitivity remains a challenge. **a**, The cumulative histogram of the amino-acid sequence lengths of a random sample of 10% of the proteins in the UniProt database of proteins. Amino-acid sequence length is reported on a log-scale along the x-axis. The solid red line indicates the median of the distribution, while the dashed blue lines indicate common context lengths of pLLMs. The yellow arrows report the context length value, and the proportion of proteins which would be truncated at that context length. **b**, The predictive performance of the ESM2-650M-based PPI inference method as a function of the amino-acid sequence lengths of the testing protein pairs. Protein pairs in the testing set are binned according to the their length, with the largest protein of each pair reported on the x-axis (Protein 1) and the smaller protein reported on the y-axis (Protein 2). The size of each circle is proportional to the number of protein pairs within each bin, and the colour of each circle reports the predictive performance (MCC@50%) when classifying pairs within the bin. The grey “x” markers denote bins for which there is no data. See Supplementary Figure 3 for visualizations of all other pLLM-based models. **c**, Stacked histograms of the proportion of correctly labelled (lighter shade) and incorrectly labelled (darker shade) testing PPI pairs for various pLLM-based PPI inference models binned according to the longest amino acid sequence in each pair (X-axis). The strict variant of SqueezeProt-SP is the only model based on a strict pLLM (red), while all other methods make use of non-strict pLLMs (blue). Each histogram reports the results for a PPI inference based on the pLLM indicated in the lower left corner. The context length of each pLLM is indicated by a dashed line. White regions indicate bins for which there is no data. **d**, Histogram showing the number of pairs for each of the bins of **(c)**, which report the longest amino acid sequence of each protein pair in the PPI testing dataset. **e**, Receiver-Operator Curve (ROC) of various PPI inference models evaluated on a testing set of human PPIs (dotted curves) and PPIs between SARS-CoV-2 and human proteins (solid curves). Each curve corresponds to one model. Six of the models are pLLM-based, while the seventh is RAPPPID (as a baseline). The SqueezeProt-SP strict and non-strict variants are coloured blue and red, respectively. See Supplementary Figure 4 and Table 2 for a detailed breakdown of model performances. **f**, Scatter plot of changes in binding free energy (y-axis) and predicted interaction probability (x-axis) between wild-type protein pairs and their mutated counterparts, as found in the SKEMPI 2.0 database. Interaction probability is inferred using the ESM2-650M-based model. For clarity of visualization circles are coloured to reflect their regional density (red is most dense and blue is least dense). The marginal plots are histograms of the distribution of changes in binding free energy (y-axis) and changes in predicted interaction probability (x-axis). See Supplementary Figure 5 and Supplementary Table 5 for a detailed breakdown for each model.

### PPI methods based on SqueezeProt-SP variants make similar predictions to those based on other pLLMs

The pLLMs analyzed in this study differ substantially from SqueezeProt-SP in their architecture, objective function, and training data. The differences between the non-pLLM based PPI inference methods and SqueezeProt-SP are even greater. We therefore investigated the concordance of predictions across protein pairs by these models in the test set to better understand if their differences are reflected therein, or if different models predict presence/absence of a PPI edge similarly.

We used the strict and non-strict variants of SqueezeProt-SP as points of reference and compared both categories of methods with them. The Cohen’s *κ* coefficient (a measure of inter-rater reliability [25, 26]) between SqueezeProt-SP models and other pLLM-based methods were similarly middling across the board (Fig. 1g, Supplementary Table 4), with an average *κ* of 0.374 (± 0.00673) and 0.326 (± 0.0115) for non-strict and strict variants, respectively. It is interesting (yet expected) to note that the pLLM-based models (that were trained on non-strict datasets) were more consistent with the non-strict SqueezeProt-SP. Non-pLLM-based methods showed less agreement with both SqueezeProt-SP variants (which is perhaps expected) in average and also showed a high variance (Supplementary Table 4).

Given that pLLM-based methods generally performed better in the PPI prediction task compared to non-pLLM approaches, controlling for their skill levels becomes important when making comparisons regarding their agreement (a requirement not satisfied by Cohen’s *κ* or other similar measures). Hence, we devised a skill-normalized concordance (SNC) measure (between 0 and 1), which accounts for any such skill-based distortions (Methods). The SNC reflected far more similarity between all methods and the two SqueezeProt-SP variants than Cohen’s *κ* (Fig. 1h, Supplementary Table 4). Despite this, a small but noticeable difference was observed, with non-pLLM-based methods showing on average a 3.70% lower degree of agreement with SqueezeProt-SP compared to pLLM-based methods. Taken together, these experiments suggest that both SqueezeProt-SP variants share a high degree of similarity with existing pLLM-based PPI inference methods. When normalizing for differences in skill, non-pLLM based methods show similar albeit lower agreement with SqueezeProt-SP. No meaningful difference between the agreement of strict and non-strict variants of SqueezeProt-SP was observed using either measures.

### Limited context-length does not impact inference of interactions between long proteins

Since the memory complexity of the attention mechanism grows quadratically as a function of sequence length, LLMs and pLLMs based on the transformer architecture must compute attention for much smaller sequences (e.g., 512) than other architectures as to make training and inference tractable. This attention cut-off is called the “context length”. Amino acid sequence lengths often surpass these limits, with 18.1% of proteins having more than 512 amino acids (some even reaching as high as 45,212 amino acids) in a randomly selected subset of UniProt (Fig. 2a) [27].

Often, these long sequences are either truncated from their C-terminus side or are omitted entirely during training, which may omit entire motifs and domains of a protein. At inference time, sequences which exceed the context length are either truncated, or windowed attention is used. It follows that pLLM-based PPI inference models might have more difficulty predicting interactions between proteins larger than the pLLM’s context-length compared to those that are smaller. To test this hypothesis, we grouped pairs of proteins in our testing dataset according to their protein length and calculated the PPI prediction performance of each group (Fig. 2b-c, Supplementary Fig. 3). Proteins larger than the context-length were truncated to the context length at inference time. It is notable that ProSE, being based on a RNN architecture, does not have context-lengths as-such.

No appreciable trend or difference in performance was observed when measuring the C3 testing performance on PPI pairs according to the length of either protein in the pair (Fig. 2b, Supplementary Fig. 3), nor when grouping pairs according to the largest of the two proteins (Fig. 2c). While our C3 PPI testing dataset contains pairs with proteins larger than 2,500 amino acids, the vast majority are below 1,000 amino acids in length (Fig. 2d). This indicates that pLLMs are seemingly capable of deriving features which are sufficient to accurately infer PPIs from the residues that fall under their context length, regardless of the length of the complete protein sequence.

### pLLM-based methods are no more capable of inferring PPIs between human and SARS-CoV-2 than other methods

Inferring interactions of proteins from organisms which differ greatly from those upon which a model is trained has long been a challenge for PPI inference models. Viral proteins are notably absent from the STRING database and are evolutionarily very distant from most other organisms present in this database [28]. Therefore, inferring the protein interactions of viral proteins is a challenging task that tests the ability of models trained on STRING DB to infer interactions that are far out of distribution from their testing set.

To systematically evaluate this, we tested pLLM-based PPI inference methods on PPIs between SARS-CoV-2 and human proteins, obtained from a previous study [29]. None of the six pLLM-based methods nor RAPPPID (a non-pLLM baseline) were capable of identifying these inter-species PPIs at rates meaningfully higher than random chance (Fig. 2e, Supplementary Fig. 4, Table 2). These results indicate that pLLM-based PPI inference models, much like their non-pLLM counterparts, struggle to make predictions on very out-of-distribution PPI interactions of viral proteins when trained on our dataset of high-confidence human PPIs derived from STRING DB.

**Table 2.** Performance of PPI models, pLLM-based and non-pLLM-based, on a testing set of interactions between SARS-CoV-2 proteins and Human proteins.

| Encoder | pLLM-based? | MCC@50% |  | Balanced Accuracy |  | AUROC |  |
| --- | --- | --- | --- | --- | --- | --- | --- |
|  |  | Average | Std. Dev. | Average | Std. Dev. | Average | Std. Dev. |
| ProtT5-XL | ✓ | $8.34 \times 10^{-5}$ | 0.00836 | 0.500 | 0.0144 | 0.507 | 0.0183 |
| ESM2-650M | ✓ | -0.0113 | 0.00349 | 0.478 | 0.00672 | 0.482 | 0.00484 |
| ProtBERT | ✓ | -0.00250 | 0.00652 | 0.495 | 0.0124 | 0.511 | 0.0126 |
| SqueezeProt-SP (Strict) | ✓ | -0.0107 | 0.00801 | 0.479 | 0.0162 | 0.468 | 0.0267 |
| SqueezeProt-SP (Non-strict) | ✓ | 0.0157 | 0.0154 | 0.530 | 0.0294 | 0.515 | 0.0302 |
| ProteinBERT | ✓ | -0.0189 | 0.00226 | 0.462 | 0.00491 | 0.444 | 0.0113 |
| ProSE | ✓ | -0.00279 | 0.00255 | 0.495 | 0.00491 | 0.488 | 0.00778 |
| RAPPPID | ✗ | -0.0133 | 0.00304 | 0.474 | 0.00630 | 0.483 | 0.00974 |

MCC@50%: Matthew’s Correlation Coefficient at a 50% threshold
AUROC: Area under the Receiver Operating Characteristic
Std. Dev.: Standard Deviation
Means and Standard Deviations are calculated for three random training/testing splits ( $n = 3$ ).

### pLLM-based PPI inference methods are not sensitive to small but important changes in amino acid sequences

The smallest discrete change in a protein sequence, namely the result of a missense DNA point mutation, can result in drastic changes in protein function, and are often complicit in cancers and hereditary diseases. Among the functional differences these mutants exhibit are alterations of their binding affinity to other proteins. Since the probabilities of PPI inference models are meant to reflect the likelihood of interaction, a reasonable expectation of a well-calibrated model is that the differences in protein interaction affinity of a mutant protein and its wild-type be reflected as a proportional change in its inferred interaction probability. We here measure the change in binding affinity as the change in Gibbs free energy (ΔΔ*G*), and the change of interaction probability as Δ*ŷ* (Methods).

Using binding affinity data of mutants and their wild-types from the SKEMPI 2.0 database [30], we calculated the Spearman correlation coefficient between Δ*ŷ* and ΔΔ*G* for both pLLM-based and non-pLLM-based methods (Fig. 2f, Supplementary Table 5). No appreciable correlation was observed as a result of these experiments. The pLLM-based PPI inference method which exhibited the correlation with the largest magnitude was ProteinBERT, with a modest Spearman coefficient of −0.122. These results suggest that the pLLM-based PPI inference models tested here are largely insensitive to small but meaningful changes in amino acid sequence.

## Discussion

Our results demonstrate that the expressive and compact representations of amino acid sequences generated by pLLMs can greatly improve machine learning models for protein analysis. As is the case with all new techniques, however, the assumptions, biases, and general consequences of incorporating pLLMs into these models warrant close scrutiny.

This study reveals a new form of data leakage in PPI inference—a task historically compromised by similar inadvertent contamination problems. Due to the innocuous nature of its source, data leakage in the PPI inference literature had long artificially inflated our understanding of the performance of these models dramatically. Due to these grave and long-lasting consequences, it is paramount for us as a community to pay close attention to possible sources of data leakage.

The particular source of data leakage we’ve identified has particular implications for the use of pre-trained pLLMs. Our study shows that pLLMs which are destined for incorporation in PPI inference models must be trained on data prepared specifically for the task. Training pLLMs from random initialized weights is a daunting task with great costs in terms of capital, time, labour, and often carbon emissions. We propose two approaches to address the costs that are most evident to us.

The first approach is to popularize the practice of creating pre-trained pLLMs which have been trained on strict datasets with held out proteins as was done in this study. The SqueezeProt-SP Strict pLLM we trained as part of this study is an example of one such model. The dataset upon which SqueezeProt-SP Strict is trained is made available for those who wish to develop and train similar “strict” models on the proteins found in SWISS-PROT, and the pipeline used to create it is outlined here in detail for those who wish to train strict variants of their models on other datasets.

The second is to develop pLLM architectures which are less costly to retrain on new datasets. Again, SqueezeProt serves as an example of a pLLM which can be trained efficiently with limited training time and computational resources while achieving good performance. Further improvements in pLLM efficiency would make it far more feasible to retrain models on datasets specifically designed for downstream tasks without risking data leakage.

Other future avenues towards addressing the risk of data leakage posed by pLLMs exist on the horizon. Further advances in machine unlearning, a process by which networks “forget” data presented at training, could have profound effects in this regard.

Importantly, while many strengths of pLLM-based PPI models have been outlined here, we also observed many challenges that remain for both pLLM-based methods and non-pLLM-based methods alike. While our previous work shows that generalization to other species is possible with traditional methods [31], generalization to species which are evolutionary very distinct from those in the training set, such as those produced by the SARS-CoV-2 virus, remain out of reach for PPI inference models of all modalities we tested. Furthermore, while PPI inference models exhibit good performance when identifying whether two proteins are likely to interact, they are not sensitive to small changes in amino acid sequences, such as those which are the result of missense point mutations, which can result in changes to the resulting protein’s binding affinity. These shortcomings do not appear to be due to the adoption of pLLMs or more traditional methods in PPI inference models, but rather a more systemic challenge posed by the current approaches to PPI inference.

Despite the challenges identified in this work, pLLMs remain tremendously promising tools for protein analysis. As is the case with most emerging technologies, it is important that practitioners and early adopters use them responsibly and with great care. As the field matures, we hope that strict training protocols will become standard practice, efficient architectures will reduce retraining costs, and new techniques will emerge to address contamination concerns.

## Methods

### Preparing C3 PPI training and testing datasets

The training and testing datasets used for downstream PPI inference tasks conformed to the C3 definition outlined by Park & Marcotte [8]. That is to say that no proteins found in the protein pairs of the testing set were found within the pairs of the training set. Similarly, no proteins which make-up the pairs of the validation set were found among the proteins of either the training or testing sets.

The PPI datasets used were previously deposited in the Zenodo repository under record number 10594149 as part of the manuscript *“INTREPPPID — an orthologue-informed quintuplet network for cross-species prediction of protein–protein interaction”* [31]. These datasets were generated using the PPI Origami software, which is also deposited in the Zenodo repository with record number 10652234 as part of the same manuscript.

The PPI Origami parameters used to generate the dataset are as follows: a confidence threshold of 90% was applied, the species of proteins in the interactions was restricted to humans by setting the taxon flag to ‘9606’. A training:testing:validation dataset size ratio of 8:1:1 was chosen. Because of the computational complexity of training pLLMs which account for the downstream PPI testing dataset, only a single random seed was chosen for the generation of the PPI dataset.

In addition to meeting the Park & Marcotte C3 definition, the dataset was processed such that the amino acid sequence of the proteins of which comprise the pairs in any given set (*i*.*e*., training, testing, or validation) were no more than 90% similar in sequence than any one protein of the other sets. Put otherwise, let 𝒫 = *P*_Tr_, *P*_Te_, *P*_V_, where *P*_Tr_, *P*_Te_, *P*_V_ are the set of proteins present in the training, testing, and validation set, respectively, such that

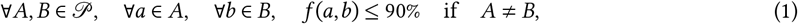

where *f* (⋅, ⋅) is a function which measures protein sequence identity.

Negative examples were generated for each split by randomly sampling pairs of proteins from the set of proteins which make up the positive examples for the same split. Put another way, the set of negative proteins for the training set is defined as

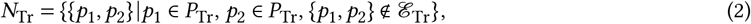

where ℰ_Tr_ is the set of all training protein pairs *E* ∈ ℰ_Tr_. Negatives were similarly generated for testing and validation sets (*N*_Te_, *N*_V_, respectively).

### Preparing “strict” and “non-strict” datasets for pLLM pre-training

Three pLLM pre-training datasets were constructed for the three pLLMs trained here: SqueezeProt-SP (strict), SqueezeProt-SP (non-strict), and SqueezeProt-U50. Both variants of SqueezeProt-SP were trained on a dataset derived from SWISS-PROT database (release number “2024_05”). The dataset for the strict variant omits any proteins that are present in the C3 PPI testing and validation sets (*P*_Te_, *P*_V_), as well as proteins which have similar sequences to them. The non-strict dataset is truncated to match the number of proteins present in the strict dataset in such a way to ensure that all the proteins omitted in the strict set are present in the non-strict set. This ensures that models trained on either the strict or the non-strict datasets have the same training data budget, and vary only in the membership of proteins present in the C3 PPI testing and validation set.

#### SqueezeProt-SP strict dataset

To ensure that all proteins in *P*_Te_ and *P*_V_, as well as proteins which are similar in sequence, are excluded from the SWISS-PROT training set, a three-pass pipeline was adopted. Proteins were considered to be similar if they shared more than 50% sequence identity. The three-pass approach was developed to ensure that, as much as possible, no similar sequences are left in the strict pre-training dataset.

First, all 572,214 protein sequences in SWISS-PROT as well as the 2,782 protein sequences from the union of *P*_Te_ and *P*_V_ were clustered using MMseqs2’s `cluster` command (version 13-45111+ds-2) [32]. Subsequently, all proteins from SWISS-PROT that share a cluster with a protein from the PPI C3 testing or validation datasets were excluded from the pLLM pre-training dataset. A total of 24,280 protein sequences were removed from SWISS-PROT using this process. The MMseqs2 parameters used with the `cluster` command were: `--max-seqs 300 --cluster-reassign 1 -s 7.5 --min-seq-id 0.5`.

To further ensure that proteins that are similar to those in the testing and validation set, a second pass was performed whereby MMseqs2’s `search` command was used to find similar proteins. MMseqs2’s `search` command works similarly to BLAST, which searches for protein sequences by performing local alignments as scoring them accordingly [33]. All matches for testing and validation proteins in the SWISS-PROT dataset as a result of the `search` command which had sequence identity greater than 50%, bit scores larger than 40, and E-values smaller than 1 were removed from the strict pre-training dataset. In this way, 14,561 protein sequences were removed from the pre-training dataset. The MMseqs2 parameters used with the `search` command were: `--num-iterations 4 -k 7 -s 5.7 -e 10000.0 --max-seqs 4000`. These values are based on those described for the “MMseqs2 profile” method described in Steinegger & Söding [32].

Finally, a third-pass, similar to the second, was performed by using the PSI-BLAST software (version 2.12.0+) to identify similar proteins in the strict pLLM pre-training datasets [33]. All matches with sequence identity greater than 50%, bit scores larger than 40, and E-values smaller than 1 were removed from the strict pre-training dataset. This resulted in 1,418 protein sequences being removed from the pre-training dataset. The PSI-BLAST parameters used were: `-num_alignments 4000 -num_iterations 4`. These values are based on those described for PSI-BLAST in Steinegger & Söding [32]. The final strict pLLM pre-training dataset contained 531,952 protein sequence; 93.0% of the size of the SWISS-PROT dataset from which it was derived.

#### SqueezeProt-SP non-strict dataset

To construct the non-strict dataset, a number of proteins from the SWISS-PROT were randomly selected to be excluded from the training set for use in the held-out evaluation datasets. The number of proteins excluded was chosen such that the number of training proteins in the non-strict dataset matches those in the strict dataset. Subsequently, proteins from SWISS-PROT were iteratively added to the training and evaluation splits according to a set of fixed conditions. Namely, if a protein sequence belongs to a protein excluded from the Strict pLLM pre-training dataset, that sequence was added to the training set. Otherwise, if the sequence did not belong to a protein in that exclusion set but had been randomly selected for the evaluation dataset at the start of the process, that sequence was assigned to the held-out evaluation set. In all other cases, the sequence was added to the training set. The final non-strict training dataset contained 531,952 protein sequences.

#### SqueezeProt-U50 dataset

Sequences from the UniRef50 dataset (release number “2024_05”) were randomly sorted between training and evaluation sets at a rate of 90% and 10%, respectively.

### SqueezeProt architecture and training procedure

All SqueezeProt variants share the same architecture, based on the SqueezeBERT model *et al*. [20]. The SqueezeBERT architecture builds on insights from the field of computer vision on how to reduce the computational complexity of convolutional neural networks (CNN) [34]. The primary way in which the SqueezeBERT architecture achieves its performance gains is by replacing all position-wise fully-connected (PFC) networks in the attention mechanism for grouped convolutions. In their study, Iandola *et al*. go on to show that “…the PFC layers of Vaswani *et al*., GPT, BERT, and similar self-attention networks can be implemented using convolutions without changing the networks’ numerical properties or behavior”[20].

As is the case for all pLLMs evaluated here, the SqueezeProt models first tokenize protein sequences on a peramino-acid basis. That is to say, each amino acid forms its own token. The SqueezeProt models were trained with absolute position embeddings and a context length of 512 tokens (equivalent to the same number of amino acids). SqueezeProt’s encoder and pooling layers have a hidden dimension of 768. Twelve such encoder layers are used, with twelve attention heads. The feed-forward layers have a size of 3,072 (also referred to as the “intermediate size”). The hidden layers are activated using the Gaussian Error Linear Unit (GELU) function and are dropped out with a 10% probability [35, 36].

SqueezeProt was trained on the masked language modelling (MLM) task (sometime referred to as a cloze or occlusion task), whereby tokens are randomly replaced with a special masking token. The network is then tasked with inferring the original tokens which the mask tokens have replced. Tokens were randomly masked with a probability of 15%. All SqueezeProt-SP variants were trained for 208 epochs, with a batch size of forty samples. The SqueezeProt-U50 model was trained for ten epochs with a batch size of sixty samples. The SqueezeProt-SP variants were each trained on a single Nvidia RTX 3090 GPU for approximately seven days. The SqueezeProt-U50 model was trained on four Nvidia RTX 3090 GPUs using distributed data parallelism over a period of approximately thirty days.

The AdamW optimizer as implemented by the HuggingFace Transformers library was used for all models trained [37]. The SqueezeProt-SP model used an initial learning rate (*α*) of 0.001, a momentum factors (*β*_1_ and *β*_2_) of 0.9 and 0.999, and zero weight decay (*λ* = 0). These correspond to the default parameters of the Transformers library’s implementation of the AdamW optimizer. A learning rate schedule which linearly decays the learning rate from its initial value of 0.001 to 0 by the last epoch is used. 16-bit mixed floating point precision was used during training. The SqueezeProt-U50 model was trained using the “1cycle” learning rate scheduler which increases a maximum learning rate of 2.7 × 10^−4^ from a minimum learning rate of 1.08 × 10^−5^ at which point it is decreased until it reaches 1.27 × 10^−9^ [38]. A cosine annealing strategy is used.

### pLLM-based PPI inference architecture and training procedure

#### Architecture

The pLLM-based PPI inference models trained here were the result of first freezing the weights of a pre-trained pLLM encoder *e* : ℝ^22×*n*^ → ℝ^*m*^, which consumes one-hot encoded amino acid sequences of length *n* and outputs an embedding of size *m*. The value of *m* is specific to the pLLM used, according to its architecture. All SqueezeProt variants, for instance, had a value of *m* equal to 768. A twin neural network was trained downstream of the encoder. The twin neural network was comprised entirely of fully-connected layers. First, an embedding projection network *g*(⋅), itself composed of two fully-connected layers *h*_0_ : ℝ^*m*^ → ℝ^32^ and *h*_1_ : ℝ^32^ → ℝ^64^, transforms the output of the pLLM encoder *z* = *e*(*x*) according to

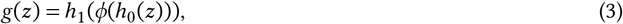

where *ϕ* is the Mish [39] non-linear activation function defined as

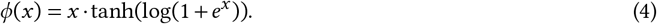

Each parameter of *h*_0_ and *h*_1_ were assigned to zero (“dropped-out”) with random probability of 15%, a regularization process known as Drop-Connect which has been shown to help avoid over-fitting [40]. For proteins *p*_0_ and *p*_1_ in a protein pair, an averaged projected latent representation *z*^′^ was computed as

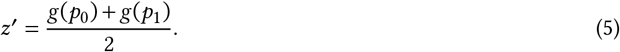

The averaged projected latent representations was then inputted into a classifier *c* : ℝ^64^ → (0, 1), whose output was the inferred probability of interaction of *p*_0_ and *p*_1_.

#### Training procedure

The network was trained for 100 epochs on the C3 training dataset described in the “*Preparing C3 PPI training and testing datasets*” section. The AdamW optimizer was used with a learning rate of 10^−3^ and weight decay of 0.02. A constant learning rate schedule was used. For each pLLM, three PPI inference models were trained, each with a different random seed. The seeds were chosen arbitrarily as the integers 1, 2, and 3.

### pLLM-based UniProt Keyword inference architecture and training procedure

### Architecture

The architecture for the UniProt Keyword prediction model was similar to that of the PPI inference model. Much like in the PPI inference model, the weights of the pre-trained SqueezeProt-SP pLLM encoder *e* : ℝ^22×*n*^ → ℝ^768^ were frozen and transform amino acid sequences of length *n* into vectors *z* = *e*(*x*). Rather than a twin neural network as in the PPI inference model, a single fully-connected network *g* : ℝ^768^ → ℝ^1086^ was trained. The output vector of *g* represents an assigned probability for each of the 1,086 keywords we aim to infer for each protein. Only keywords assigned to more than 10 proteins are included in this analysis. The network *g* was constructed of three hidden layers *h*_0_ : ℝ^768^ → ℝ^256^, *h*_1_ : ℝ^256^ → ℝ^512^, and *h*_2_ : ℝ^512^ → ℝ^1086^. Thus, we obtained the inferred probabilities for each UniProt keyword of a protein *ŷ* ∈ (0, 1)^1086^ from its one-hot encoded sequence *x* ∈ ℝ^22×*n*^ composed of *n* amino acids by the network defined as

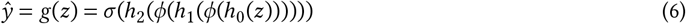

where *ϕ* is the Mish non-linear activation function and *σ* is the sigmoid function.

#### Training

The keyword inference task is a multi-label task with a high-degree of imbalance between labels. Furthermore, proteins are sparsely labelled, with 0.7% of keywords being assigned to each protein on average. To help address these challenges, we adopted the asymmetric loss (ASL) function for multi-label tasks described by Ridnik *et al*. [41]. The ASL function is similar to Lin *et al*.’s focal loss, but introduces asymmetric focusing parameters *γ* ^+^ (the positive focusing hyper-parameter) and *γ* ^−^ (the negative focusing hyper-parameter) [42]. ASL also introduces “shifted probabilities” *ŷ*_*m*_ for negative samples, which involves the thresholding of negative samples with very low probabilities (*i*.*e*., *ŷ*_*m*_ = max(*ŷ* −*m*, 0) where *m* is a tunable parameter, and *ŷ* are inferred probabilities).

We modified the ASL function so as to re-weight classes inversely to their frequencies. We applied the following loss weights *w*_*c*_ ∈ *W* for each class *c* ∈ {0,…, 1085}. Loss weights for each class are defined as

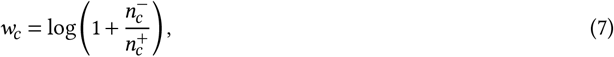

where 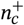 and 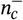 are the number of proteins which are labelled with class *c*, and those which are not, respectively. This results in a final weight vector *W* = [*w*_0_,…, *w*_1085_]. Applying class weights to the final loss function greatly helped mitigate the degradation in performance we observed due to severe class imbalance. The final loss function was composed of a positive (*L*_+_) and negative loss (*L*_−_) defined as

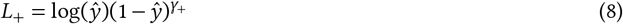

and

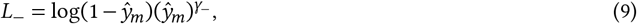

respectively. These loss functions were then combined to form a final loss function

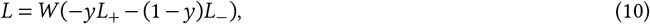

where *y* are the ground-truth labels. For our purposes, we used the following hyperparamter values: *γ*_+_ = 1, *γ*_−_ = 4, and *m* = 0.05. The network was trained for 20 epochs with a batch size of 100. The network was optimized with 16-bit mixed precision with the Adam optimizer on a RTX 3090 GPU.

### Skill-Normalized Concordance

Most measures that exist for assessing the inter-rater concordance of different methods (*e*.*g*., Cohen’s or Fleiss’s *κ*) do not account for the relative skill of the raters (their ability to correctly make predictions). As a result, the difference in skill between a high-skill rater and a low-skill rater will influence their concordance scores. This is not desirable in cases where one intends on understanding agreement between methods independent of their relative skill. To address this, we devised a method for measuring concordance among different classifiers while normalizing for their relative skills.

Consider two classification methods with differing skill. The model with the better skill is labelled as model *b* and that with the worse skill is labelled as *w*. More concretely,

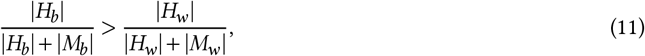

where *H*_*b*_ and *H*_*w*_ are the set of the indices of the correctly classified testing samples (“hits”) of models *b* and *w*, respectively. *M*_*b*_ and *M*_*w*_ are the set of incorrectly classified testing samples (“misses”) of models *b* and *w*, respectively. The skill-normalized concordance (SNC) function *λ* is defined as

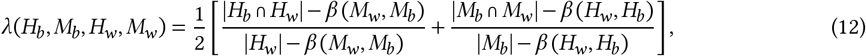

where for two sets of indices *R* and *S*,

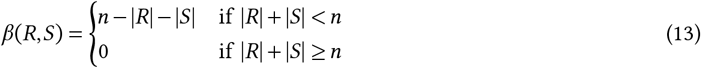

and *n* is the total number of testing samples (*i*.*e*., *n* = |*H*_*w*_ |+|*M*_*w*_ | = |*H*_*b*_ |+|*M*_*b*_|). The range of SNC is between 0 and 1, with higher scores indicating higher agreement between methods (while controlling for their skills).

To better understand the utility of SNC compared to other concordance measures, consider four hypothetical binary predictors *A, B, C*, and *X*. When inferring the same testing set consisting of 10 samples with known ground-truth labels (*y* ∈ {0, 1}^10^), the result for each method can be represented as the sets

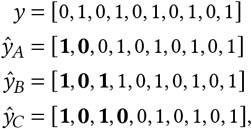

where *y* is the ground-truth and *ŷ*_*A*_, *ŷ*_*B*_, and *ŷ*_*C*_ are the resulting predicted labels from models *A, B*, and *C*, respectively, with incorrectly labelled elements being marked in bold face. From these results, we can see that these methods have differing skill, with *A, B*, and *C* having accuracies of 80%, 70%, and 60%, respectively. Despite these differences, it is immediately apparent that these methods have the highest possible agreements, given their accuracies.

The pairwise Cohen’s *κ* coefficients for these methods are equal to *κ*(*ŷ*_*A*_, *ŷ*_*B*_) = 0.8, *κ*(*ŷ*_*A*_, *ŷ*_*C*_) = 0.6, and *κ*(*ŷ*_*B*_, *ŷ*_*C*_) = 0.8. As a result, the high agreement of these classifiers (given their differing skill) is not reflected in these scores. On the other hand, the SNC assigns a value of one to any two pairs of methods above, reflecting that these methods have the highest possible agreement with each other (given that their skill is different).

SNC similarly accounts for skill when methods largely disagree. Consider a model *X*, which is used to infer the labels of the same testing dataset as *A, B*, and *C* with predictions

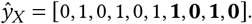

Model *X* disagrees maximally with *A, B*, and *C* for a method that scores 60% accuracy on the dataset in question. This is indeed reflected in the skill-normalized concordance between *ŷ*_*X*_ and *ŷ*_*A*_, *ŷ*_*B*_, and *ŷ*_*C*_, which is zero in all cases. Cohen’s *κ* reports agreements between the same methods ranging between −0.19 and −0.6, once more reflecting the relative skill of the methods compared.

While Cohen’s *κ* coefficient remains a more relevant measure of inter-rater agreement in instances where random-chance agreement is of greater import or true labels are missing, SNC offers complimentary insight into the commonality between raters independent of their skill.

### Evaluating PPI inference performance on SARS-CoV-2 networks

The protein-protein interaction network between SARS-CoV-2 viral proteins and human proteins expressed in HEK-293T were collected from a previous study [29]. Information of these inter-species interactions comes from affinity-purification mass spectrometry data which describe the PPIs of all but three SARS-CoV-2 viral proteins after being expressed in human HEK-293T cells. Protein pairs were labelled as interacting if the pair had a MiST score greater than or equal to 0.7, a SAINTexpress Bayesian false-discovery rate (BFDR) less than or equal to 0.05, and an average spectral count greater than or equal to 2. These thresholds were chosen to match those used in the original study [29]. All other pairs were labelled as not interacting.

### Evaluating the impact of mutations on PPI inference performance

Data on binding affinity was curated, processed, and made available by Strokach *et al*. [43]. From this dataset, only data from the SKEMPI database (labelled as “skempi++”), which report the ΔΔ*G* as the result of mutations, was considered. Two-sided Spearman correlation was calculated using the SciPy library [44].

### Choice of evaluation metrics

No single evaluation metric is sufficient to adequately summarize every aspect of performance for a given model in every context. For this reason, we have included multiple performance metrics where possible in our analysis. The performance of binary classification tasks, such as the PPI inference task, are reported using metrics which report performance for one (MCC, *F* −1, Accuracy) or a range of threshold values (AUROC, AP). Care is taken to ensure that chosen metrics are resilient to class imbalance (*e*.*g*, MCC and Balanced Accuracy) when evaluation sets are not balanced. For ranking tasks, such as the keyword annotation task, relevant ranking metrics were chosen (*e*.*g*., NDCG, LRAP, Coverage Error, Top-*k* Accuracy).

## Data Availability

All datasets generated for the analyses herein, as well as data produced by those analyses, are made available under a the Creative Commons Attribution-NonCommercial-ShareAlike 4.0 International license. Data and can be accessed via GitHub: https://github.com/Emad-COMBINE-lab/pllm-ppi-data-leakage.

## Code Availability

All code used to conduct the analyses and create the figures herein are made available under the GNU Affero General Public License v3. The code-base can be accessed via GitHub: https://github.com/Emad-COMBINE-lab/pllm-ppi-data-leakage.

## Acknowledgements

This work was supported by grants from Natural Sciences and Engineering Research Council of Canada (NSERC) [RGPIN-2019-04460] (A.E.) and Canada Foundation for Innovation (CFI) JELF [project 40781] (A.E.). This research was enabled in part by support provided by Calcul Québec (www.calculquebec.ca) and the Digital Research Alliance of Canada (alliancecan.ca)

## Author contributions

A.E. and J.S. contributed to project conceptualization, methodology as well as writing, editing, and reviewing. A.E. contributed to supervision and funding acquisition. J.S. contributed to software, data curation, investigation and visualization. All authors read and approved the final article.

## Notes

https://github.com/Emad-COMBINE-lab/pllm-ppi-data-leakage

https://emad-combine-lab.github.io/pllm-ppi-data-leakage/

### Competing Interest Statement

The authors have declared no competing interest.

### Summary of Updates

The manuscript is reformatted, and insights are discussed in more details.

