## Supplementary File for "A flaw in using pre-trained pLLMs in protein-protein interaction inference models"

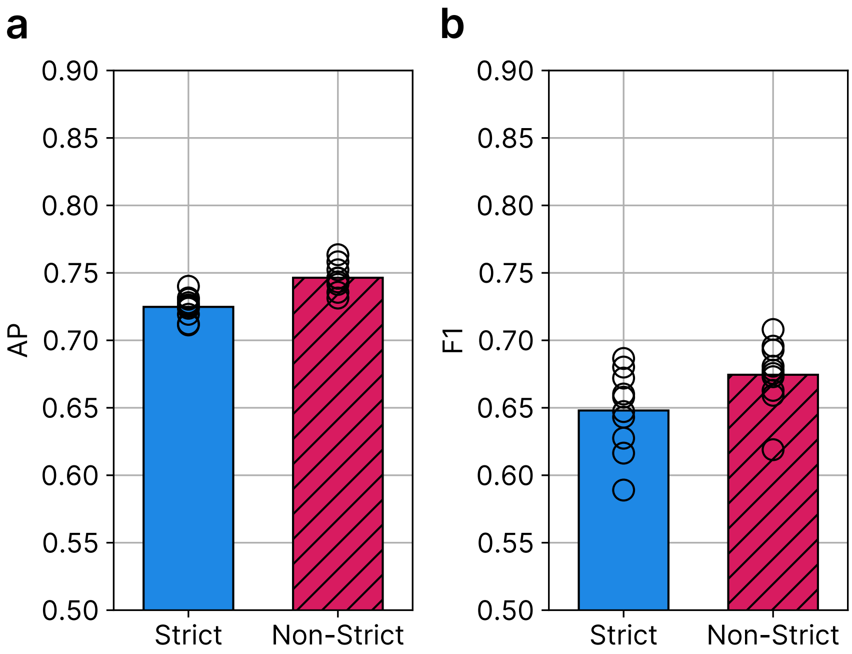


**Supplementary Figure 1:** The mean performance of the PPI inference models on the PPI testing set based on strict (blue) and non-strict (red, hatched) SqueezeProt-SP pLLM encoders, as measured by AP **(a)** and $F_1$ score **(b)**. The mean AP of the strict and non-strict models is 0.725 (±0.0083) and 0.746 (±0.00918) respectively, and the mean $F_1$ score of the strict and non-strict models is 0.648 (±0.0288) and 0.674 (±0.0233). Mean and standard deviation are calculated for the results of ten models trained identically but for different random seeds used to initialize weights and sample training examples. Markers indicate the performance of each of the ten models.

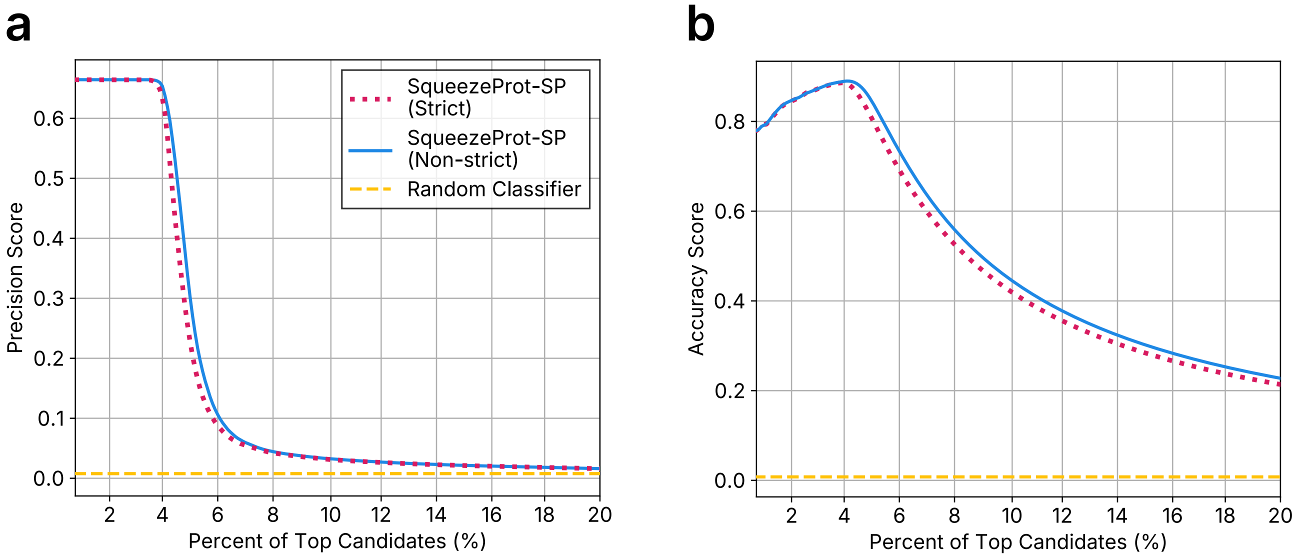


**Supplementary Figure 2:** Overall testing performance, as measured by precision score **(a)** and accuracy score **(b)**, among the top ranked candidates of strict (solid blue) and non-strict (dotted red) pLLM-based keyword models. The performance of a random classifier (dashed yellow) is also included for comparison.

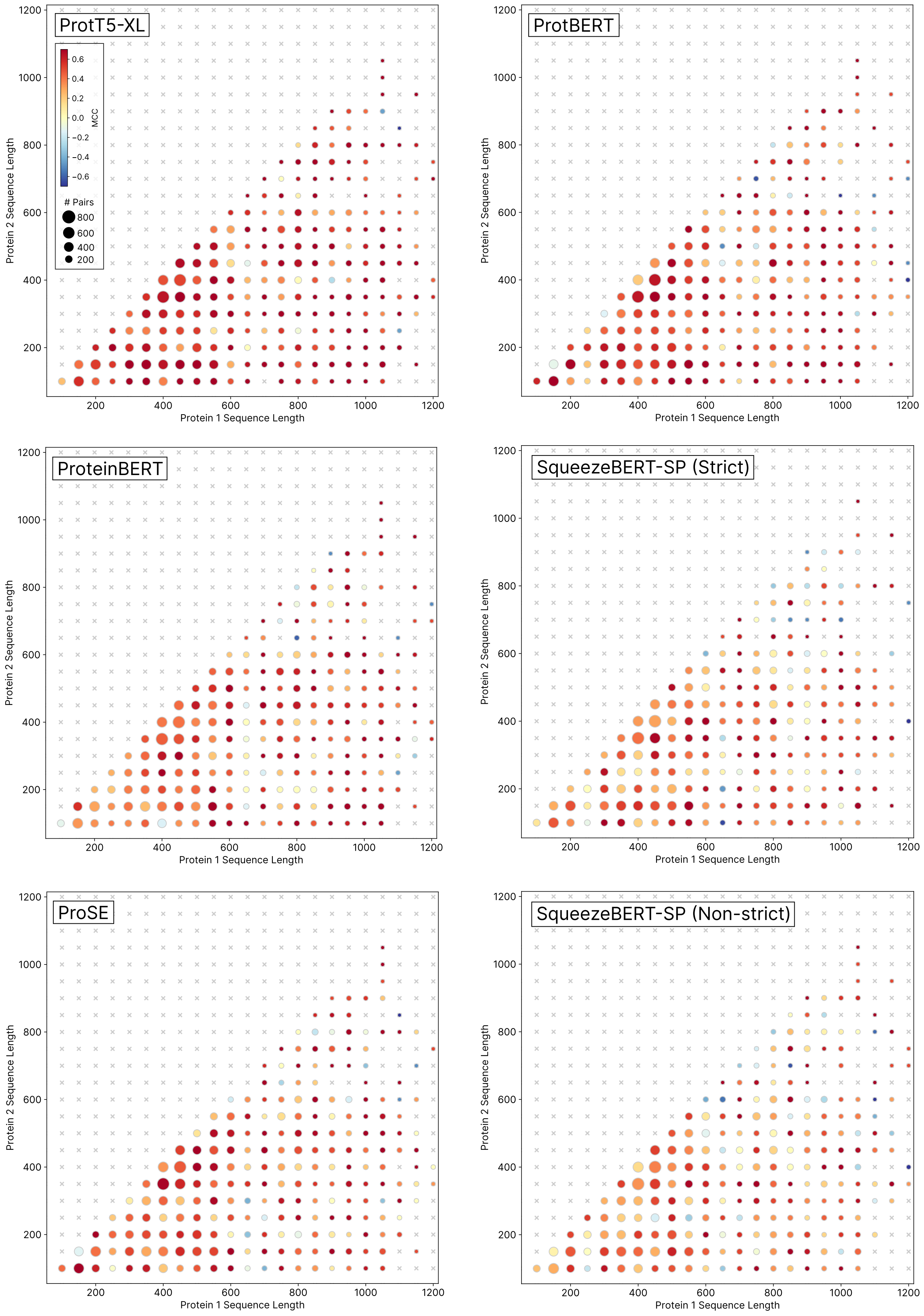


**Supplementary Figure 3:** (*Caption on next page.*)

**Supplementary Figure 3:** (***Previous page.***) Heat maps of the predictive performance of various pLLMs according to the amino-acid sequence length of the proteins in the testing pairs. The model name in the upper-left corner indicate the pLLM used in the pLLM-based PPI inference model for each analysis. Protein pairs in the testing set are binned according the their length, with the largest protein of each pair reported on the X-axis (Protein 1) and the smaller protein reported on the Y-axis (Protein 2). The size of each circle is proportional to the number of protein pairs within each bin, and the the colour of each circle reports the testing performance of the model indicated when classifying pairs within the bin, as reported by MCC. The grey "X" markers denote bins for which there is no data.

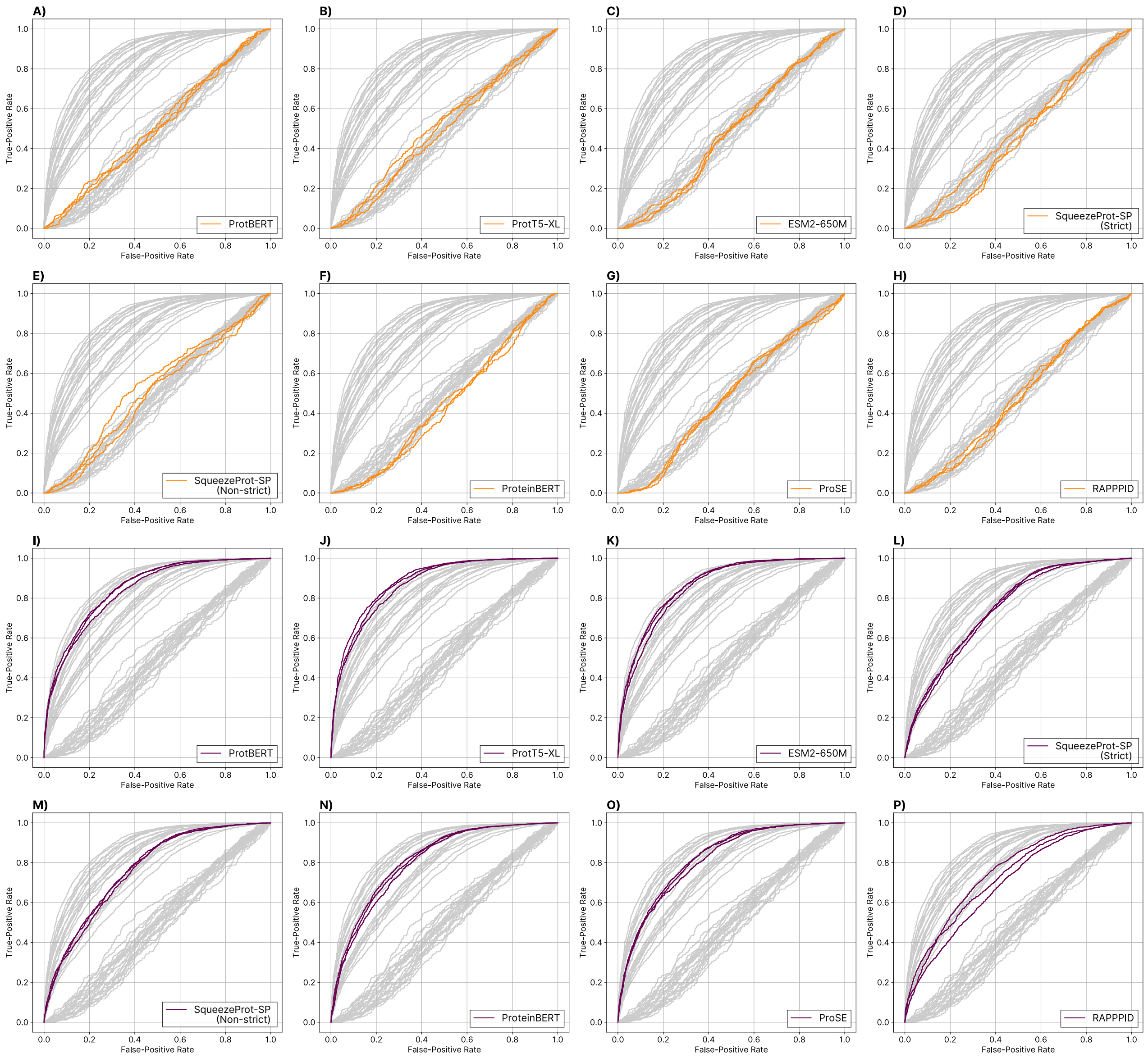


**Supplementary Figure 4:** Receiver-Operator Curve (ROC) of various PPI inference models evaluated on a testing set of human PPI pairs (purple curves) and PPIs between SARS-CoV-2 and human proteins (orange curves). Each plot reports the result of the testing ROCs of pLLM-based PPI inference models, with the pLLM indicated in its lower-left corner. Results are also reported for one non-pLLM PPI inference model (RAPPPID). Three coloured ROCs per plot each represent the results for a model trained using different random seed. Light grey ROCs in each plot report all other methods.

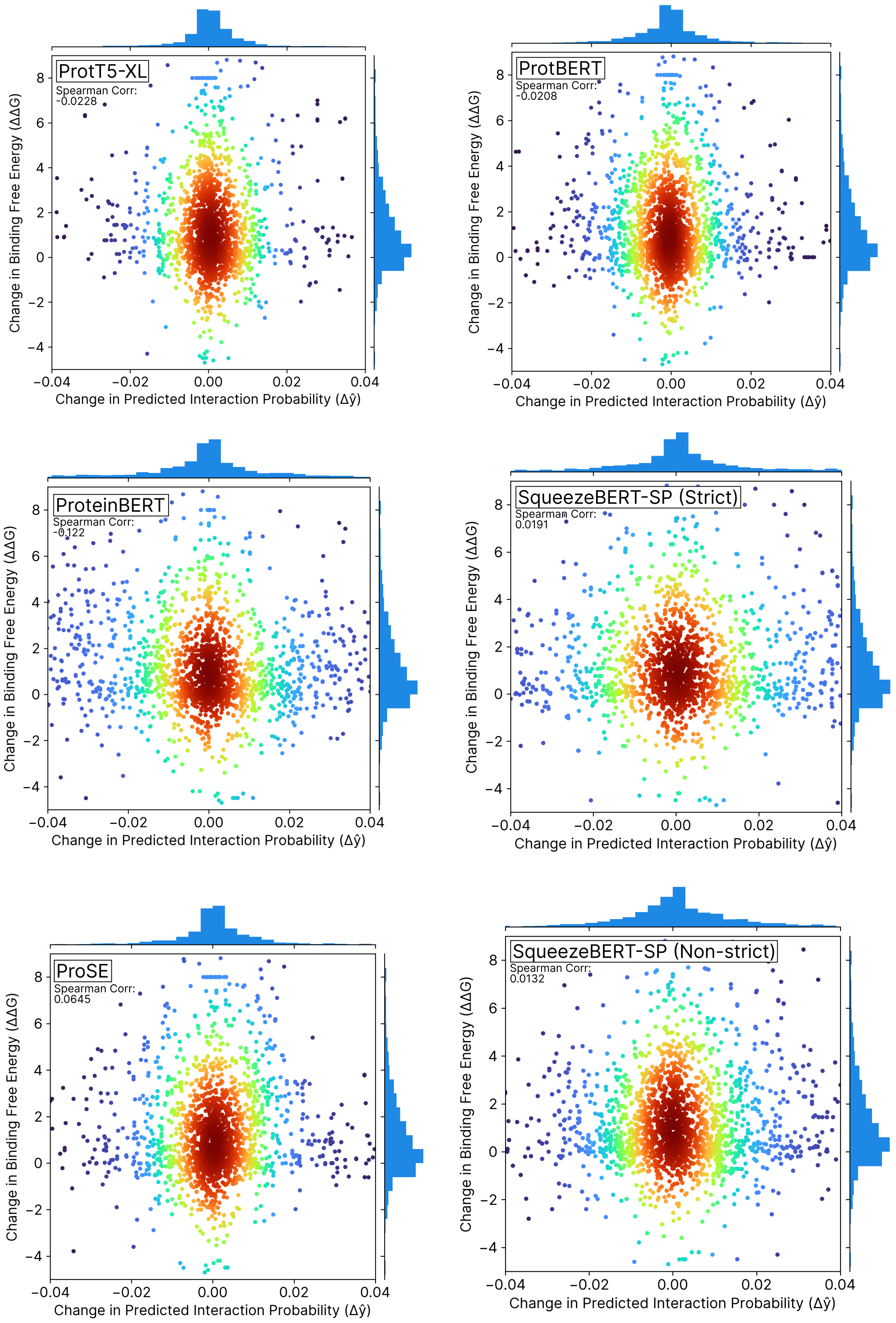


**Supplementary Figure 5:** (*Caption on next page.*)

**Supplementary Figure 5: (*Previous page.*)** Scatter plots of changes in binding free energy (y-axis) and predicted interaction probability (x-axis) between wild-type protein pairs and their mutated counterparts, as found in the SKEMPI 2.0 database. Interaction probability is inferred using a pLLM-based PPI inference method, with the pLLM indicated in the upper-left corner. The colour of points in the plot indicate density (red is most dense, blue is least dense). The marginal plots are histograms of the distribution of changes in binding free energy (y-axis) and changes in predicted interaction probability (x-axis). The Spearman correlation of the changes in binding free energy and changes in predicted interaction probability is reported in the upper-left corner of each plot.

**Supplementary Table 1:** Characteristics of pLLMs studied herein.

| pLLM Name | Year | # Params ($\times 10^7$) | Context Length | Training Tasks | Training Dataset | # Training Proteins ($\times 10^6$) | Tokens per Second (inference) |
|---|---|---|---|---|---|---|---|
| ESM2-650M | 2023 [10] | 65.0 | 1,024 | MLM | UniRef50 [9] | 66 | 6,417.6 |
| ProteinBERT | 2022 [2] | 1.60 | 1,024 | MLM, Protein Annotation | UniRef90 [9] GO [4] | 194 | 8,047.6 |
| ProtBERT | 2021 [6] | 42.0 | 2,048 | MLM | UniRef100 [9] | 420 | 7,716.4 |
| ProtT5-XL | 2021 [6] | 40.9 | 512 | MLM | UniRef50 [9] | 66 | 4,928.0 |
| ProSE | 2021 [1] | 5.9 | n.a. | MLM, Contact Map Prediction, Structure Similarity | UniRef90 [9] SCOPe [7, 3] | 194 | 2,849.4 |
| SqueezeProt-SP | This Study | 2.83 | 512 | MLM | SWISS-PROT† [5] | 0.5 | 39,559.4 |
| SqueezeProt-U50 | This Study | 2.83 | 512 | MLM | UniRef50 [9] | 66 | 39,559.4 |

MLM: Masked Language Model, GO: Gene Ontology

†The strict variant omits proteins that are present in downstream PPI testing datasets, while the non-strict variant includes them.

**Supplementary Table 2:** PPI inference performance on a testing set for both strict and non-strict variants of SqueezeProt-SP

| | MCC@50% | | AUROC | | AP | | $F$-1 Score | | Accuracy | |
|---|---|---|---|---|---|---|---|---|---|---|
| Seed | Non-Strict | Strict | Non-strict | Strict | Non-strict | Strict | Non-strict | Strict | Non-strict | Strict |
| 1 | 0.360 | 0.305 | 0.769 | 0.744 | 0.741 | 0.711 | 0.659 | 0.616 | 0.679 | 0.650 |
| 2 | 0.363 | 0.341 | 0.768 | 0.748 | 0.746 | 0.724 | 0.663 | 0.658 | 0.680 | 0.670 |
| 3 | 0.327 | 0.346 | 0.758 | 0.753 | 0.731 | 0.725 | 0.619 | 0.672 | 0.660 | 0.673 |
| 4 | 0.372 | 0.341 | 0.770 | 0.756 | 0.746 | 0.730 | 0.675 | 0.643 | 0.685 | 0.669 |
| 5 | 0.380 | 0.365 | 0.767 | 0.759 | 0.736 | 0.740 | 0.693 | 0.680 | 0.690 | 0.682 |
| 6 | 0.355 | 0.339 | 0.768 | 0.752 | 0.743 | 0.728 | 0.673 | 0.660 | 0.677 | 0.669 |
| 7 | 0.381 | 0.342 | 0.775 | 0.755 | 0.746 | 0.731 | 0.678 | 0.647 | 0.690 | 0.670 |
| 8 | 0.384 | 0.361 | 0.780 | 0.755 | 0.758 | 0.726 | 0.681 | 0.687 | 0.691 | 0.681 |
| 9 | 0.409 | 0.297 | 0.787 | 0.737 | 0.763 | 0.712 | 0.708 | 0.589 | 0.704 | 0.643 |
| 10 | 0.388 | 0.316 | 0.780 | 0.741 | 0.752 | 0.719 | 0.696 | 0.627 | 0.694 | 0.656 |
| Mean | 0.372 | 0.335 | 0.772 | 0.750 | 0.746 | 0.725 | 0.674 | 0.648 | 0.685 | 0.666 |
| St. Dev. | 0.0221 | 0.0224 | 0.00853 | 0.00713 | 0.0968 | 0.00875 | 0.0245 | 0.0303 | 0.0120 | 0.0126 |

MCC@50%: Matthew's Correlation Coefficient at a 50% threshold.
AUROC: Area under the Receiver Operating Characteristic.
AP: Average Precision.
Std. Dev.: Standard Deviation.

**Supplementary Table 3:** Mean keyword model performance by category and metric.

| Keyword Category | Model | Mean NDCG▲ | Mean LRAP▲ | Mean Coverage Error▼ | Mean Top-3 Accuracy▲ | Mean Top-10 Accuracy▲ |
|---|---|---|---|---|---|---|
| Cellular component | SqueezeProt-SP (Strict) | 0.541 (±7.76E-3) | 0.588 (±3.01E-3) | 13.1 (±2.11) | 0.742 (±6.64E-4) | 0.742 (±1.28E-2) |
| | SqueezeProt-SP (Non-Strict) | 0.547 (±7.33E-3) | 0.596 (±3.75E-3) | 13.1 (±2.13) | 0.742 (±1.92E-4) | 0.765 (±1.21E-2) |
| | Random Model | 0.188 (±7.56E-4) | 0.214 (±5.27E-4) | 77.8 (±6.03E-1) | 0.0116 (±2.88E-4) | 0.0115 (±1.45E-4) |
| Ligand | SqueezeProt-SP (Strict) | 0.348 (±3.36E-4) | 0.739 (±1.51E-4) | 5.45 (±7.67E-1) | 0.762 (±6.22E-4) | 0.835 (±1.69-2) |
| | SqueezeProt-SP (Non-Strict) | 0.348 (±4.88E-4) | 0.738 (±9.14E-5) | 5.47 (±1.15) | 0.762 (±1.03E-4) | 0.875 (±1.06E-2) |
| | Random Model | 0.156 (±7.89E-4) | 0.518 (±9.08E-4) | 24.4 (±3.64E-1) | 0.0177 (±2.69E-4) | 0.0174 (±4.17E-4) |
| PTM | SqueezeProt-SP (Strict) | 0.317 (±2.45E-3) | 0.825 (±4.58E-4) | 3.29 (±4.10E-1) | 0.836 (±5.07-4) | 0.357 (±2.10E-2) |
| | SqueezeProt-SP (Non-Strict) | 0.318 (±3.14E-3) | 0.825 5.93E-4 | 3.30 (±4.70E-1) | 0.836 (±1.51E-4) | 0.362 (±8.53E-3) |
| | Random Model | 0.145 (±1.01E-3) | 0.620 (±6.64E-4) | 12.3 (±3.12E-1) | 0.0223 (±4.93E-4) | 0.0222 (±1.84E-4) |
| Molecular Function | SqueezeProt-SP (Strict) | 0.290 (±8.25E-4) | 0.363 (±1.07E-2) | 39.2 (±6.31E-2) | 0.928 (±1.11E-16) | 0.835 (±2.03E-2) |
| | SqueezeProt-SP (Non-Strict) | 0.291 (±4.33E-4) | 0.364 9.78E-3 | 39.3 (±9.71E-2) | 0.928 (±1.11E-16) | 0.872 (±7.96E-3) |
| | Random Model | 0.161 (±4.85E-4) | 0.249 (±6.41E-4) | 92.2 (±6.54E-2) | 0.0077 (±7.39E-4) | 0.00768 (±2.91E-4) |
| Biological Function | SqueezeProt-SP (Strict) | 0.267 (±9.06E-4) | 0.542 (±1.21E-3) | 58.4 (±4.28E-2) | 0.897 (±1.11E-16) | 0.840 (±1.15E-2) |
| | SqueezeProt-SP (Non-Strict) | 0.266 (±7.88E-4) | 0.542 1.57E-3 | 59.0 (±4.29E-2) | 0.897 (±1.11E-16) | 0.893 (±3.54E-3) |
| | Random Model | 0.112 (±4.03E-4) | 0.398 (±3.93E-4) | 1.86E2 (±6.77E-2) | 0.00309 (±3.60-4) | 0.00311 (±3.21E-4) |
| Domain | SqueezeProt-SP (Strict) | 0.233 (±1.15E-4) | 0.797 (±6.07E-4) | 2.77 (±1.87E-2) | 0.881 (±1.26E-4) | 0.446 (±3.52E-3) |
| | SqueezeProt-SP (Non-Strict) | 0.232 (±6.02E-5) | 0.797 4.00E-4 | 2.79 (±2.57E-2) | 0.882 (±3.42E-4) | 0.451 (±3.88E-3) |
| | Random Model | 0.131 (±8.00E-4) | 0.671 (±6.62E-4) | 8.11 (±3.25E-2) | 0.0203 (±7.42E-4) | 0.021 (±4.33E-4) |

LRAP: Label Ranking Average Precision [11], NDCG: Normalized Discounted Cumulative Gain [8]
▲Higher values indicate better performance.
▼Lower values indicate better performance.
Standard deviation reported in brackets.

**Supplementary Table 4:** Concordance measures between SqueezeProt variants and other PPI inference methods.

| | | SqueezeProt-SP (Strict) | | SqueezeProt-SP (Non-strict) | |
|---|---|---|---|---|---|
| Method | pLLM-based? | Cohen's $\kappa$ | SNC | Cohen's $\kappa$ | SNC |
| ESM2-650M | ✓ | 0.386 | 0.533 | 0.343 | 0.551 |
| ProtT5 | ✓ | 0.373 | 0.521 | 0.317 | 0.524 |
| ProtBERT | ✓ | 0.369 | 0.508 | 0.329 | 0.523 |
| ProSE | ✓ | 0.371 | 0.507 | 0.314 | 0.504 |
| ProteinBERT | ✓ | 0.370 | 0.510 | 0.328 | 0.523 |
| | Mean | 0.374 | 0.516 | 0.326 | 0.525 |
| | Std. Dev. | 0.00698 | 0.0111 | 0.0115 | 0.0168 |
| RAPPPID | ✗ | 0.311 | 0.502 | 0.301 | 0.526 |
| D-SCRIPT | ✗ | 0.215 | 0.426 | 0.195 | 0.445 |
| Richoux *et al.* | ✗ | 0.112 | 0.551 | 0.0765 | 0.549 |
| SPRINT | ✗ | 0.00452 | 0.442 | 0.0128 | 0.424 |
| PIPR | ✗ | -0.00792 | 0.563 | -0.00220 | 0.584 |
| | Mean | 0.127 | 0.497 | 0.117 | 0.506 |
| | Std. Dev. | 0.137 | 0.0620 | 0.129 | 0.0685 |

SNC: Skill-Normalized Concordance
Std. Dev.: Standard Deviation.

**Supplementary Table 5:** Correlation between the difference of the inferred interaction probability ($\Delta\hat{y}$) and Gibbs Free Energy ($\Delta\Delta G$).

| Encoder | pLLM-based? | Spearman Correlation |
|---|---|---|
| ProtT5-XL | ✓ | -0.0228 |
| ESM2-650M | ✓ | -0.0539 |
| ProtBERT | ✓ | -0.0208 |
| SqueezeProt-U50 | ✓ | -0.0766 |
| ProteinBERT | ✓ | -0.122 |
| ProSE | ✓ | 0.0645 |
| RAPPPID | ✗ | -0.0369 |

[4] The Gene Ontology Consortium, Suzi A Aleksander, James Balhoff, Seth Carbon, J Michael Cherry, Harold J Drabkin, Dustin Ebert, Marc Feuermann, Pascale Gaudet, Nomi L Harris, David P Hill, Raymond Lee, Huaiyu Mi, Sierra Moxon, Christopher J Mungall, Anushya Muruganugan, Tremayne Mushayahama, Paul W Sternberg, Paul D Thomas, Kimberly Van Auken, Jolene Ramsey, Deborah A Siegele, Rex L Chisholm, Petra Fey, Maria Cristina Aspromonte, Maria Victoria Nugnes, Federica Quaglia, Silvio Tosatto, Michelle Giglio, Suvarna Nadendla, Giulia Antonazzo, Helen Attrill, Gil dos Santos, Steven Marygold, Victor Strelets, Christopher J Tabone, Jim Thurmond, Pinglei Zhou, Saadullah H Ahmed, Praoparn Asanitthong, Diana Luna Buitrago, Meltem N Erdol, Matthew C Gage, Mohamed Ali Kadhum, Kan Yan Chloe Li, Miao Long, Aleksandra Michalak, Angeline Pesala, Armalya Pritazahra, Shirin C C Saverimuttu, Renzhi Su, Kate E Thurlow, Ruth C Lovering, Colin Logie, Snezhana Oliferenko, Judith Blake, Karen Christie, Lori Corbani, Mary E Dolan, Harold J Drabkin, David P Hill, Li Ni, Dmitry Sitnikov, Cynthia Smith, Alayne Cuzick, James Seager, Laurel Cooper, Justin Elser, Pankaj Jaiswal, Parul Gupta, Pankaj Jaiswal, Sushma Naithani, Manuel Lera-Ramirez, Kim Rutherford, Valerie Wood, Jeffrey L De Pons, Melinda R Dwinell, G Thomas Hayman, Mary L Kaldunski, Anne E Kwitek, Stanley J F Laulederkind, Marek A Tutaj, Mahima Vedi, Shur-Jen Wang, Peter D'Eustachio, Lucila Aimo, Kristian Axelsen, Alan Bridge, Nevila Hyka-Nouspikel, Anne Morgat, Suzi A Aleksander, J Michael Cherry, Stacia R Engel, Kalpana Karra, Stuart R Miyasato, Robert S Nash, Marek S Skrzypek, Shuai Weng, Edith D Wong, Erika Bakker, Tanya Z Berardini, Leonore Reiser, Andrea Auchincloss, Kristian Axelsen, Ghislaine Argoud-Puy, Marie-Claude Blatter, Emmanuel Boutet, Lionel Breuza, Alan Bridge, Cristina Casals-Casas, Elisabeth Coudert, Anne Estreicher, Maria Livia Famiglietti, Marc Feuermann, Arnaud Gos, Nadine Gruaz-Gumowski, Chantal Hulo, Nevila Hyka-Nouspikel, Florence Jungo, Philippe Le Mercier, Damien Lieberherr, Patrick Masson, Anne Morgat, Ivo Pedruzzi, Lucille Pourcel, Sylvain Poux, Catherine Rivoire, Shyamala Sundaram, Alex Bateman, Emily Bowler-Barnett, Hema Bye-A-Jee, Paul Denny, Alexandr Ignatchenko, Rizwan Ishtiaq, Antonia Lock, Yvonne Lussi, Michele Magrane, Maria J Martin, Sandra Orchard, Pedro Raposo, Elena Speretta, Nidhi Tyagi, Kate Warner, Rossana Zaru, Alexander D Diehl, Raymond Lee, Juancarlos Chan, Stavros Diamantakis, Daniela Raciti, Magdalena Zarowiecki, Malcolm Fisher, Christina James-Zorn, Virgilio Ponferrada, Aaron Zorn, Sridhar Ramachandran, Leyla Ruzicka, and Monte Westerfield. The gene ontology knowledgebase in 2023. *Genetics*, 224(1):iyad031, May 2023.

[5] The UniProt Consortium, Alex Bateman, Maria-Jesus Martin, Sandra Orchard, Michele Magrane, Shadab Ahmad, Emanuele Alpi, Emily H Bowler-Barnett, Ramona Britto, Hema Bye-A-Jee, Austra Cukura, Paul Denny, Tunca Dogan, ThankGod Ebenezer, Jun Fan, Penelope Garmiri, Leonardo Jose Da Costa Gonzales, Emma Hatton-Ellis, Abdulrahman Hussein, Alexandr Ignatchenko, Giuseppe Insana, Rizwan Ishtiaq, Vishal Joshi, Dushyanth Jyothi, Swaathi Kandasaamy, Antonia Lock, Aurelien Luciani, Marija Lugaric, Jie Luo, Yvonne Lussi, Alistair MacDougall, Fabio Madeira, Mahdi Mahmoudy, Alok Mishra, Katie Moulang, Andrew Nightingale, Sangya Pundir, Guoying Qi, Shriya Raj, Pedro Raposo, Daniel L Rice, Rabie Saidi, Rafael Santos, Elena Speretta, James Stephenson, Prabhat Totoo, Edward Turner, Nidhi Tyagi, Preethi Vasudev, Kate Warner, Xavier Watkins, Rossana Zaru, Hermann Zellner, Alan J Bridge, Lucila Aimo, Ghislaine Argoud-Puy, Andrea H Auchincloss, Kristian B Axelsen, Parit Bansal, Delphine Baratin, Teresa M Batista Neto, Marie-Claude Blatter, Jerven T Bolleman, Emmanuel Boutet, Lionel Breuza, Blanca Cabrera Gil, Cristina Casals-Casas, Kamal Chikh Echioukh, Elisabeth Coudert, Beatrice Cuche, Edouard De Castro, Anne Estreicher, Maria L Famiglietti, Marc Feuermann, Elisabeth Gasteiger, Pascale Gaudet, Sebastien Gehant, Vivienne Gerritsen, Arnaud Gos, Nadine Gruaz, Chantal Hulo, Nevila Hyka-Nouspikel, Florence Jungo, Arnaud Kerhornou, Philippe Le Mercier, Damien Lieberherr, Patrick Masson, Anne Morgat, Venkatesh Muthukrishnan, Salvo Paesano, Ivo Pedruzzi, Sandrine Pilbout, Lucille Pourcel, Sylvain Poux, Monica Pozzato, Manuela Pruess, Nicole Redaschi, Catherine Rivoire, Christian J A Sigrist, Karin Sonesson, Shyamala Sundaram, Cathy H Wu, Cecilia N Arighi, Leslie Arminski, Chuming Chen, Yongxing Chen, Hongzhan Huang, Kati Laiho, Peter McGarvey, Darren A Natale, Karen Ross, C R Vinayaka, Qinghua Wang, Yuqi Wang, and Jian Zhang. Uniprot: the universal protein knowledgebase in 2023. *Nucleic Acids Research*, 51(D1):D523–D531, January 2023.

[6] Ahmed Elnaggar, Michael Heinzinger, Christian Dallago, Ghalia Rehawi, Yu Wang, Llion Jones, Tom Gibbs, Tamas Feher, Christoph Angerer, Martin Steinegger, Debsindhu Bhowmik, and Burkhard Rost. Prottrans: Toward understanding the language of life through self-supervised learning. *IEEE Transactions on Pattern Analysis and Machine Intelligence*, 44(10):7112–7127, October 2022.

[7] Naomi K. Fox, Steven E. Brenner, and John-Marc Chandonia. Scope: Structural classification of proteins–extended, integrating scop and astral data and classification of new structures. *Nucleic Acids Research*, 42(Database issue):D304–309, January 2014.

[8] Kalervo Järvelin and Jaana Kekäläinen. Cumulated gain-based evaluation of ir techniques. 20:422–446, October 2002.

[9] Baris E. Suzek, Yuqi Wang, Hongzhan Huang, Peter B. McGarvey, Cathy H. Wu, and the UniProt Consortium. Uniref clusters: a comprehensive and scalable alternative for improving sequence similarity searches. *Bioinformatics*, 31(6):926–932, March 2015.

[10] Robert Verkuil, Ori Kabeli, Yilun Du, Basile I. M. Wicky, Lukas F. Milles, Justas Dauparas, David Baker, Sergey Ovchinnikov, Tom Sercu, and Alexander Rives. Language models generalize beyond natural proteins. December 2022.

[11] Xi-Zhu Wu and Zhi-Hua Zhou. A unified view of multi-label performance measuresa unified view of multi-label performance measures. (arXiv:1609.00288), September 2016. arXiv:1609.00288 [cs.LG].
